# Do auditory deviants evoke cortical state changes under anaesthesia? A proof of concept study

**DOI:** 10.1101/2024.12.11.627934

**Authors:** Laura H Bohórquez, Manuel S Malmierca, Adam Hockley

## Abstract

Context-dependent sensory processing within the predictive coding framework relies on detecting mismatches between incoming stimuli and internal predictive models. Sensory deviants elicit prediction errors, seen as enhanced neural responses, that update these models and influence attention and behaviour. Although prediction errors have been widely observed across brain regions, the downstream processes remain poorly understood. In this study, we recorded electrocorticography in 5 urethane-anaesthetised rats and identified cortical slow oscillations, characterised by spontaneous transitions between “Up” and “Down” states. Deviant stimuli in an auditory oddball paradigm evoked an initial positive prediction error, followed by a prolonged, all-or-nothing response which spread in a travelling wave across the cortex. Identified as putative evoked cortical Up states, these responses were not evoked by standards, omissions or a many-standards control. Up states following deviants occurred more recently after a previous Up state when compared to spontaneous Up states. In preliminary data from an awake rat, long-latency Up states were not present spontaneously or evoked. In a different rat, anaesthetic depth was key to spontaneous and evoked Up states, with more robust Up/Down states and more reliable triggering of Up states under deeper anaesthesia. These results suggest that sensory deviants may cause shifts in cortical state under anaesthesia, explaining the long-latency MMR under anaesthesia and sleep.

**Highlights:**

- **What is the central question of this study?** Whether long-latency deviant-evoked mismatch responses could be produced by deviant stimuli in the oddball paradigm evoking Up states.
- **What is the main finding and its importance?** In anaesthetised rat, responses to unexpected stimuli show two components, an early ‘mismatch negativity’ followed by an ‘Up’ state, that may be related to deviant activation of cortical ensembles by unexpected stimuli.

## INTRODUCTION

Context modulates neural processing of sensory stimuli. The brain suppresses responses to predictable inputs and amplifies responses to unexpected ones. This deviance detection mechanism enhances the salience of unpredictable events that often carry survival-relevant information. Predictive coding provides a framework for understanding contextual processing, wherein the brain (1) encodes regular patterns to form internal predictive models, (2) detects deviations from those models, and (3) updates the models to guide attention and behavioural choices. Extensive work has characterised prediction error responses to deviant auditory stimuli across a range of paradigms (Audette et al., 2022; Dürschmid et al., 2016; Gelens et al., 2024; Parras et al., 2017; Shymkiv et al., 2025). However, the downstream mechanisms following prediction errors remain less well understood. In monkeys, auditory cortex neurons generate early (∼100 ms) and late (∼300 ms) responses to deviant stimuli (Gong et al., 2024). The early response temporally coincides with the mismatch negativity (MMN), while the late response correlates with behavioural decisions. Changes to attention and behavioural choice can be produced by cortical state changes (Harris & Thiele, 2011; Poulet & Crochet, 2019; Speed et al., 2019), and prediction errors can induce cognitive state transitions between behavioural tasks (Cole et al., 2024a). Because cortical state changes support attentive decision-making (Crochet & Petersen, 2006; Greenberg et al., 2008; Poulet & Crochet, 2019; Poulet & Petersen, 2008), prediction errors may initiate these transitions to direct attention and shape behavioural responses.

Cortical slow oscillations exemplify cortical state changes and emerge during the synchronised brain states of slow-wave sleep and anaesthesia. These oscillations occur globally across the cortex, thalamus, and hippocampus, and involve reliable transitions between “Up” states of high neural activity and “Down” states of low activity (Adamantidis et al., 2019; Neske, 2016; Sanchez-Vives, 2020; Staresina et al., 2015). During slow-wave sleep and deep anaesthesia, the brain generates Up/Down transitions at a frequency of 0.2–0.5 Hz (Steriade et al., 1993a). Studies using methods ranging from in vitro single-cell recordings to scalp electroencephalography consistently record slow oscillations (Maria V. Sanchez-Vives & David A. McCormick, 2000; Steriade et al., 1993b).

Thalamic input plays a critical role in initiating cortical Up states. Although the cortex continues to generate slow oscillations after deafferentation (Timofeev et al., 2000), inactivating the thalamus markedly reduces their frequency (David et al., 2013; Lemieux et al., 2014; Rigas & Castro-Alamancos, 2007). Thalamocortical activity consistently precedes the onset of Up states, indicating that this input likely initiates the transitions and regulates their frequency (Contreras & Steriade, 1995). External somatosensory and visual stimuli also trigger cortical Up states, with each wave originating in the corresponding sensory region (Hasenstaub et al., 2007; Ji & Wilson, 2007; Petersen et al., 2003). Based on this evidence, we propose the hypothesis that deviant stimuli in the oddball paradigm may increase thalamocortical drive and thereby evoke Up states.

We selected urethane anaesthesia to model cortical state transitions because it induces the full spectrum of the sleep cycle, including regular, rhythmic alternations between Up and Down states (Clement et al., 2008; Durán et al., 2021; Gretenkord et al., 2017, 2020; Jercog et al., 2017; Mondino et al., 2024a; Pagliardini et al., 2013). We recorded electrocorticography (ECoG) signals under urethane and observed spontaneous Up states during silent periods. Unexpected stimuli in the oddball paradigm consistently evoked Up states. The binary nature of Up state triggering generated bimodal amplitude distributions, and the resulting all-or-nothing, long-latency responses matched the profile of cortical slow oscillations. Deviant stimuli in auditory cortex elicited an initial peak at 50 ms and 20 µV, followed by a later peak at 200 ms and 200 µV, and these responses propagated across cortical areas as traveling Up state waves. The timing of preceding Up states constrained the likelihood of new ones, consistent with a refractory period. Anaesthetic depth shaped the brain’s responsiveness to deviant stimuli, determining whether they triggered an Up state.

## METHODS

### Ethical approval

All animal experimental procedures were performed under protocols established by the Directive of the European Communities (86/609/CEE, 2003/65/CE, and 2010/63/UE) and RD53/2013 Spanish Legislation for the use and care of animals. All the details of the study were approved by the Bioethics Committee of the University of Salamanca (USAL-ID-887). Seven Female Long-Evans rats were obtained from Janvier Labs (Le Genest-Saint-Isle, France), and experiments were performed on animals weighing 210-280g. Rats were housed on a 12/12 h light-dark cycle, with food and water available *ad libitum*. At termination of experiments, animals were killed by perfusion or pentobarbital overdose and decapitated, following Annex IV in the European Directive 2010/63/EU. The authors understand and comply with the ethical principles of The Journal of Physiology.

### Anaesthetic induction and auditory brainstem responses

All experiments were made inside a chamber with acoustic insulation and electrical protection. For acute experiments, anaesthesia was induced with urethane (1.9 g kg^−1^, i.p.; Sigma-Aldrich, St Louis, U.S.A.), then a further 0.315 - 0.63 g kg^−1^ dose at 45 minutes if required to induce surgical anaesthetic depth. This dose is greater than that used previously to induce robust synchronised states consisting of Up and Down states (Durán et al., 2021). For chronic implant anaesthesia was induced with ketamine (Ketamidor, 50 mg kg^-1^, i.p.; Richter Pharma, Wels, Austria) and xylazine (Rompun, 5 mg kg^-1^, i.p.; Bayer, Berlin, Germany) and recovery protocol was followed as previously (Hockley et al., 2025). Corneal and rear paw withdrawal reflexes were routinely monitored to assess anaesthetic depth. Ophthalmic gel (Lipolasic 2 mg/g; Bausch Lomb, Vaughan, Canada) was applied for avoid dry eyes in the animals. During all experimental procedures, the animals were artificially ventilated through a tracheotomy with O_2_, monitoring the CO_2_ levels; except for the chronic implanted animal, where no tracheotomy was performed. Temperature was maintained at 36 - 38°C using a rectal probe and a homeostatic heating blanket (Cibertec, Madrid, Spain) placed under the animal.

Sound stimuli were presented using Tucker-Davis Technologies (TDT; Alachua, FL, USA) hardware and presented monaurally to the right ear through a closed system using a custom-made earphone (calibrated to ensure a flat spectrum; ±2 dB at 0.5 - 44 kHz) coupled to a custom-made cone, as a substitute for traditional ear bars. Initially, auditory brainstem responses (ABRs) were recorded to check for normal hearing in animals. Three needle electrodes were placed into the skin, one at the dorsal midline close to the neural crest, one behind the left pinna, and one behind the right pinna. ABRs were recorded in response to tone bursts (4, 8, 16 and 32 kHz; 5 ms duration, 1 ms rise/fall times, 21 Hz presentation rate, 512 repetitions in 10 dB steps from 0–80 dB SPL; Tucker-Davis Technologies RZ6 & Medusa 4Z preamp). ABR thresholds were analysed with TDT BioSigRZ and all animals showed thresholds ≤40 dB SPL and were considered normal hearing for the following experiments.

### Surgery and ECoG array implantation

Following ABRs, dexamethasone (Cortexonavet, 0.25 mg kg−1, i.m.; Syva, León, Spain) and atropine (0.1 mg kg−1, s.c.; Braun, Rubi, Spain) were administered to reduce cerebral oedema and bronchial secretions, respectively. Lidocaine was also injected around the pinna tissue to achieve a higher anaesthetic level in these areas. The ears were prepared by removing pinna cartilage to obtain easier access the ear canal, and a tracheotomy allowed artificial ventilation. The animal was placed in the stereotactic frame with the upper jaw attached to a bite bar and the hollow ear bars into the ear canals. The speaker was placed into the right ear bar, while the hollow left ear bar was filled with plasticine. A midline incision was made along the midline of the head, from 5 mm rostral of bregma to the nuchal ridge, as well as small lateral incisions (∼3 mm) at each end. Connective tissue was removed and the temporalis muscle moved to allow access to AC-L, AC-R, mPFC-L and mPFC-R. An electrode array was produced, made of 8 silver wires (0.125 mm diameter; WPI, Sarasota, USA) heated at one end to form a ball, to prevent damage to the dura mater, and attached to a ZCA-EIB16 (TDT, Alachua, USA) ZIF-clip connector board (Berger et al., 2018). A total of 10 burr holes were drilled for electrode placement and anchor screws, at the locations shown in Fig 1A. Silver ball electrodes were placed in the burr holes on the rostral and caudal AC and mPFC. After electrode insertion, the drill holes were filled with Kwik-Cast silicone sealant (World Precision Instruments, Hitchin, UK) and then the entire array covered with dental acrylic (Duralay; Reliance Dental Manufacturing; Alsip, IL, USA). Finally, skin was sutured around the implant. 6 Animals were not allowed to recover, and first recordings were made approximately 1 hour after the end of the surgery. One rat was chronically implanted, after the surgery, the animal was treated during the following 3-5 days until the disappearance of the symptoms. This included the administration of anti-inflammatory every 24h (Metacam, 1 mg kg^−1^, s.c.; Boehringer Ingelheim, Ingelheim am Rhein, Germany), analgesic every 12h (Bupaq 0.1 mg kg^−1^, s.c.; Richter Pharma AG, Wels, Austria) and a topical triantibiotic every 24h (Dermisone; Novartis, Basilea, Switzerland).

**FIGURE 1:**
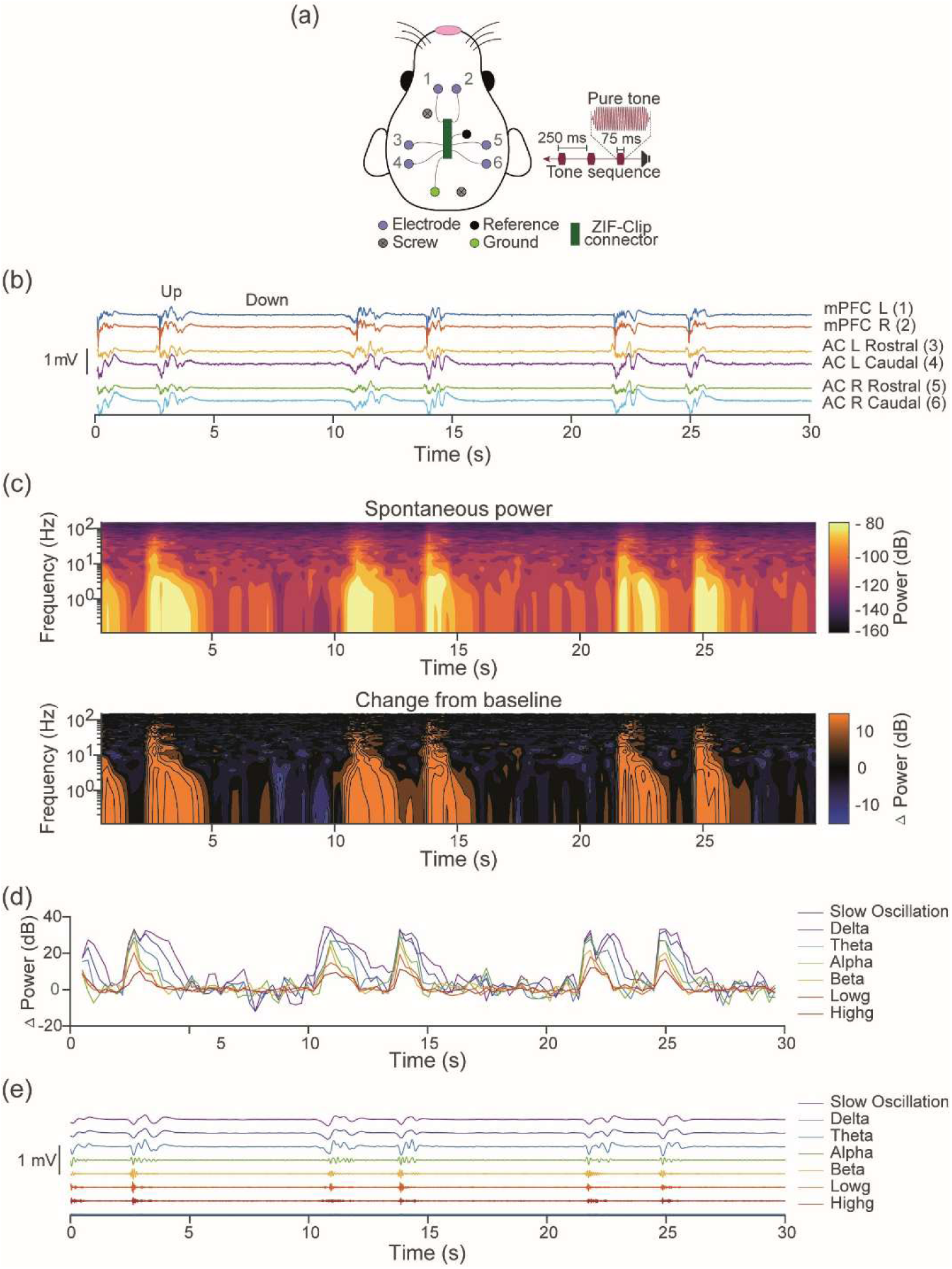
Spontaneous Up and Down states in the urethane-anaesthetised rat. **(a)** Schematic of the six recording sites, two over mPFC (1 & 2) and four over AC, bilaterally placed with rostral (3 & 5) and caudal sites (4 & 6). **(b)** Example simultaneous LFP traces from all six electrode sites show that Up states are coincidental across electrode sites. Data from 30s of silence is shown, during which cortical Up and Down states are observed. **(c)** Spontaneous cortical Up states are associated with increased activity across oscillatory bands. Top shows spontaneous power for the LFP trace in B. Spontaneous increases occur across traditional oscillators bands and align with LFP trace peaks. Bottom shows change in power from baseline. **(d)** Oscillatory power traces demonstrate coincidental increases across all power bands. **(e)** Low- and high-frequency components of the LFP are shown by filtering for specific frequency bands.

### Neural recording and analysis

Custom MATLAB (Mathworks, Natick, USA) code was used to generate stimuli patterns and control presentation through TDT Synapse software. Stimuli were output from the TDT RZ6 and presented through a speaker to the right ear. A TDT ZC16 headstage was used to connect the implanted electrodes to a TDT PZ5 preamp connected to a TDT RZ2 processor. The data were filtered (low-pass 7 kHz) and recorded using TDT Synapse software at a sampling rate of 3 kHz. We analysed the TDT stream data as local field potential (LFP) data using custom Matlab scripts. The stream was downsampled to 763 Hz, then filtered using an 8th order bandpass filter (0.1-1 Hz for LFPs; 0.1-150 Hz for oscillation analysis). Oscillatory analysis bands used were: delta, 2-4 Hz; theta 4-8 Hz; alpha 8-14 Hz; beta 14-30 Hz; low gamma 30-70 Hz; high gamma 70-150 Hz.

### Quantification and statistical analysis

Up state initiation found in LFP traces using the Matlab findpeaks function (MinPeakDistance = 382; MinPeakProminence = 3 standard deviations). Peaks that had another peak in the preceding 2s were removed, as they were deemed to be later waves of the same Up state. Identified Up state onsets peaks were 200-300 µV, larger than traditional evoked responses of 20-50 µV (Jung et al., 2013a). Up states were identified as deviant-evoked if they occurred 100-500ms after a deviant (DEV). All further analysis, statistical testing and plotting of all data were performed using custom MATLAB scripts. Cross-correlation synchrony was calculated using a bin size of 50 ms. Total synchrony was quantified as the total synchrony measurement for a 250 ms window centered at the peak synchrony timing. Synchrony measures for were statistically compared using two-sample t-tests, comparing the synchrony of the DEV-Up to the synchrony of the standard (STD)-Up for the standard preceding the DEV.

## RESULTS

### Urethane anaesthesia induces spontaneous Up and Down states

We surgically implanted ECoG arrays in five rats, placing electrodes over left and right auditory cortex (AC-L and AC-R) and medial prefrontal cortex (mPFC-L and mPFC-R) (Figure 1a), and began recordings 4–6 hours after anaesthetic induction. In the absence of sensory stimulation, we observed spontaneous Up states occurring simultaneously across all six electrodes at a frequency of approximately 0.2 Hz in filtered (0.1–150 Hz) LFPs Figure 1b). These Up states featured increased activity lasting 1–3 seconds across traditional oscillatory bands, with high-frequency activity nested within the Up state (Figure 1c–e). These findings confirm that neural activity under urethane anaesthesia consists of spontaneous Up and Down state dynamics.

### Auditory deviants evoke cortical Up states

To determine whether the auditory oddball paradigm evokes cortical Up states, we presented a repeating tone of 10 kHz interspersed with rare DEV tones of 14.142 kHz (75 ms duration, 250 ms stimulus onset asynchrony (SOA), 6.45% DEV probability, inter-DEV interval 3–4.75 s, 60 dB SPL) while recording ECoG activity as shown in Figure 1. The standard (STD) preceding the DEV was used for analyses. The inter-DEV interval produced a deviant presentation rate of approximately 0.2–0.35 Hz, which varied randomly but matched the spontaneous Up state frequency. Up states can be identified in LFPs (Gretenkord et al., 2017; Torao-Angosto et al., 2021) thus following previous methods using LFPs we identified likely Up state initiations as the first negative peaks in the filtered (0.1–10 Hz) LFP traces from the mPFC-L electrode (Figure 2a, black dots). We selected mPFC-L for Up state detection due to its superior signal-to-noise ratio, although Up states appeared simultaneously across all electrode sites, as shown in Figure 1b.

**FIGURE 2:**
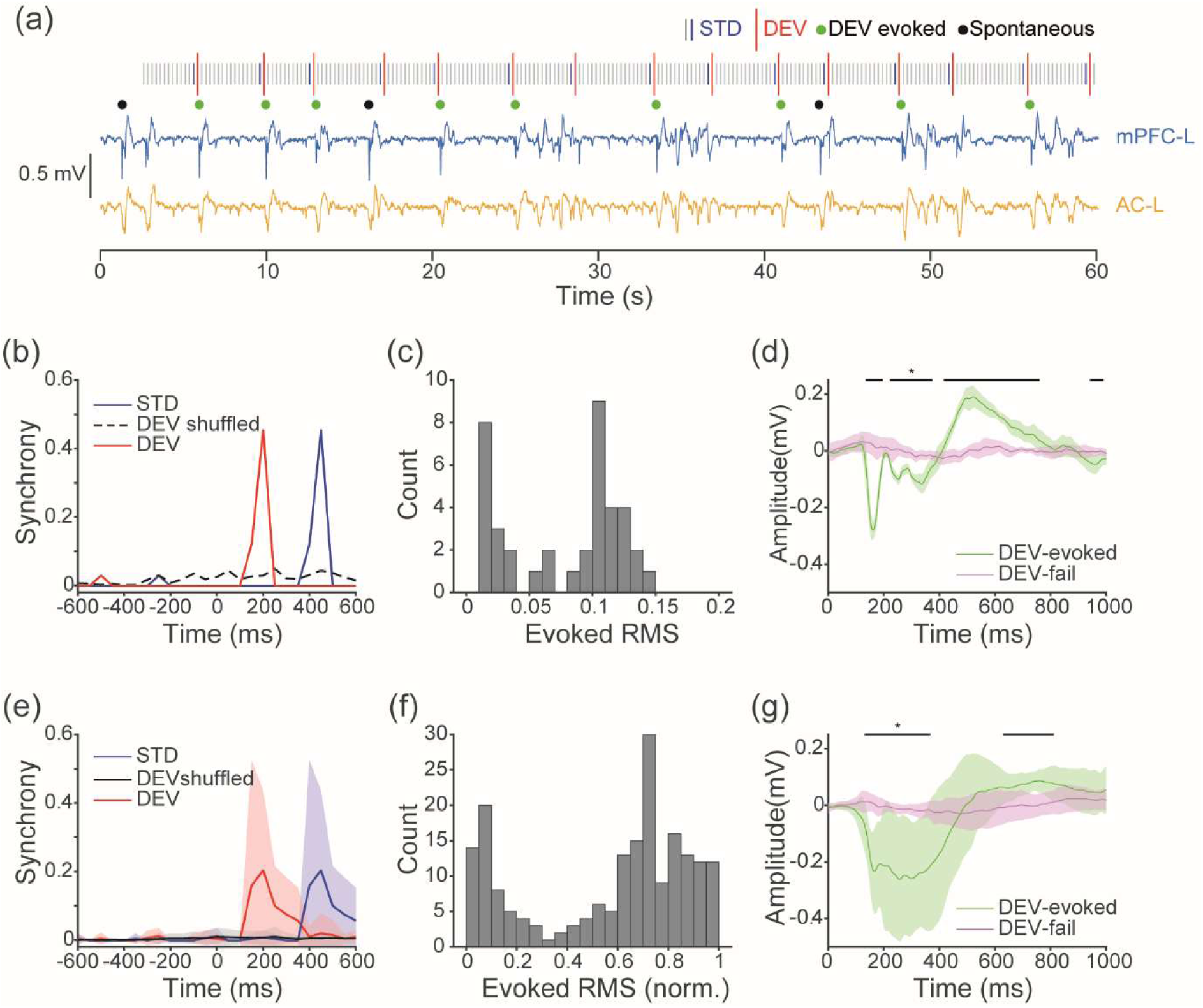
Auditory deviants initiate Up states in the mPFC. **(a)** LFP traces from the mPFC-L (blue) and AC-L (yellow), electrode sites 1 and 3, respectively from an example animal. An auditory oddball paradigm was presented with a SOA of 250 ms and inter-DEV interval of 3 - 4.75 s (11 – 18 STDs (blue bars; pseudorandomised). Up state initiations occurred either spontaneously (black dots) or following auditory DEVs (DEV red bar; Evoked Ups green dots). For clarity only the initial 30s are shown, full stimuli length was 620 stimuli, 40 DEVs, 155s. **(b)** Cross-correlation synchrony between Up state initiation and DEV or STD stimuli presentation for the example animal in panel A. Synchrony in the DEV condition is observed with a ∼200ms latency, demonstrating reliable initiation of cortical Up state by the auditory DEV. In comparison, such synchrony was not present in a DEV time shuffled condition, or in response to the STD. STDs were followed by a DEV 250ms later, producing the ‘STD’ synchrony response at ∼450ms. **(c)** Histogram of evoked RMS in the 0-500ms window following DEV stimuli, for the example animal in panel A. Response RMS to DEV stimuli is bimodal, indicating an all-or-nothing response, not a simple auditory response. **(d)** For the example animal in panel A, we partitioned mPFC LFP responses to DEV stimuli by whether an Up state is initiated. When DEVs evoked an Up state (green), a negative peak occurred at ∼200 ms then a long-latency response up to 850 ms in duration. In comparison, when DEVs failed to evoke an Up state (purple), no other response is observed. Black bar mark significance between DEV-evoked and DEV-fail (two-sample t-test, α=0.05). 0 ms represents the DEV stimulus onset. Mean ± 95% CI. **(e-g)** The same analyses as panels B-D but mean responses across all trials for each animal were calculated before population analyses (*n*=5).

In all animals, a high proportion of DEV stimuli were followed by cortical Up states (M = 57%, SD = 20.1). These responses frequently lasted 1–3 seconds, indicating sustained changes in cortical activity. In quantifying the reliability of this effect, we assessed the temporal relationship between stimulus onset and Up state initiation using cross-correlation synchrony. We compared DEV-Up synchrony to the synchrony between Up states and the STD tones that occurred immediately before each deviant. To confirm that the results were not due to phase alignment or entrainment, we have used a time-shuffled control of deviant data, where deviant onset times were shuffled (iterations = 100; −1 to 1s; 100ms intervals). Across animals, DEV stimuli consistently evoked Up states, with DEV-Up synchrony significantly exceeding STD-Up synchrony and peaking around 200 ms after stimulus onset (two-sample t-test; t(8) = 2.9677, p = 0.0179 Figure 2e). DEV-Up synchrony also significantly exceeding that of the time-shuffled condition (two-sample t-test; t(8) = 2.8626, p = 0.0211).

To test whether deviant responses followed a bimodal distribution, consistent with the all-or-nothing nature of the possible Up state initiation, we calculated the root-mean-square (RMS) amplitude from 0 to 500 ms following each DEV and plotted the distribution. This analysis revealed a clear bimodal response profile in both the example animal (Figure 2c) and in the group average after normalizing RMS values within each animal (Figure 2f). We statistically confirmed a non-normal distribution by z-score standardizing the RMS values then comparing to a standard normal distribution with a one-sample Kolmogorov-Smirnov test (p = <0.0001). We confirmed a bimodal distribution, with a skewness of −0.5509, excess kurtosis of −1.1507 and a bimodal coefficient of 0.6940 (threshold for bimodality is 0.555)(Pfister et al., 2013). We then separated the deviant responses by the presence or absence of an Up state initiation. When an Up state occurred, the signal exhibited a negative peak at approximately 200 ms, followed by a long-latency component that extended up to 850 ms (example trace in Figure 2d, animal means in Figure 2g). RMS between 100-400ms of DEV-evoked Up states were significantly greater than DEV-fail responses, tested using a linear mixed-effect model with DEV/Fail as a fixed effect and Animal and Trial as random effects (ANOVA; F(1, 188) = 300.95, p < 0.0001).

We next tested a classical oddball paradigm with a 500 ms SOA, 10% DEV probability, and a mean inter-DEV interval of 5 s, using a flip-flop design with 10 and 14.142 kHz tones (75 ms duration, 60 dB SPL; Figure 3a). As in Figure 2, we observed spontaneous and deviant-evoked responses that followed an all-or-nothing, long-latency profile. To investigate suitable sites of Up state initiation and propagation, we compared deviant responses across AC-L, AC-R, and mPFC-L. When a deviant could triggered an Up state, response amplitudes exceeded those observed during non-evoked trials (Figure 3b,e). The AC-L showed the shortest latency response to deviants (45.87 ms), followed by later responses in AC-R (186.11 ms) and mPFC-L (263.43 ms). We confirmed the latency difference using a linear mixed-effect model, with electrode location as a fixed effect and Animal as a random effect (ANOVA; F(1, 10) = 36.623, p=0.000119). Cohen’s d effect sizes and confidence intervals for the 5 tested animals were 1.140, 95% CI [-0.134, 2.359]; 1.056, 95% CI [-0.200, 2.258]; 1.693, 95% CI [0.275, 3.045]; 1.4096, 95% CI [0.069, 2.687]; 1.724, 95% CI [0.297, 3.084].

**FIGURE 3:**
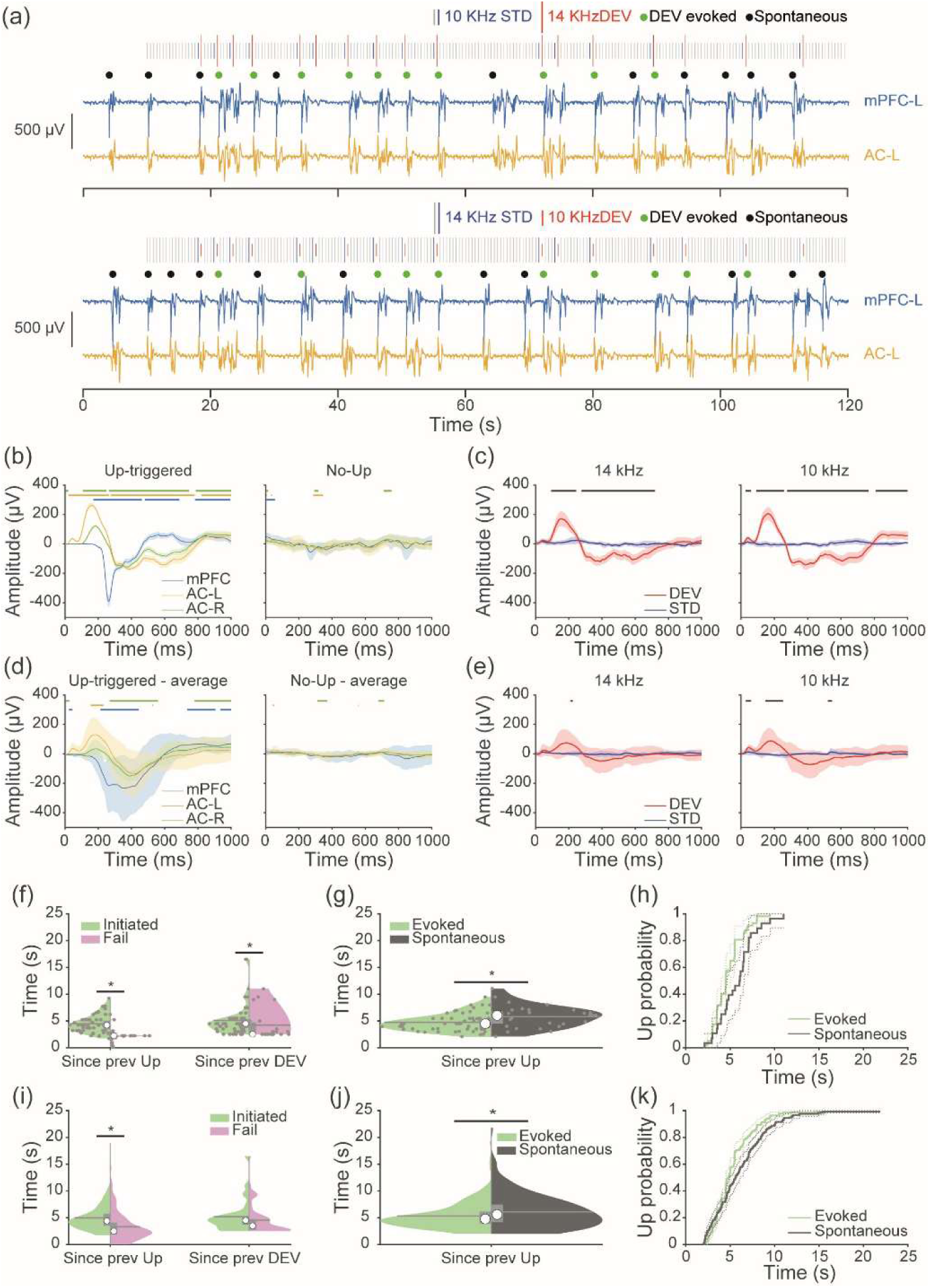
A generic auditory Oddball paradigm also initiates cortical Up states. **(a**) LFP traces from mPFC-L (blue) and AC-L (yellow) electrodes for an example animal. For each animal, a classical oddball paradigm was presented with a 500 ms SOA and 10% DEV probability. Stimuli length was 75 ms, 10% DEV probability, 82 minutes. Up state initiations were identified and classified as either evoked (PFC response occurring 100-500ms after a DEV; green dots) or spontaneous (black dots). **(b)** Mean ± 95% CI plots of DEV response from an example animal at different electrodes sites when the DEV is followed by an Up state (left) or not followed by an Up state (right). Green, yellow and blue bars indicate the presence of significance from 0 for AC-R, AC-L and mPFC-L, respectively (one sample t-test, t(58) *P*<0.05). (**c)** Comparison of LFP responses to DEV and STD stimuli of 14 kHz (left) and 10kHz (right) in the AC-L from an example animal. Mean ± 95% CI. Black bar indicates the significance (two sample unpaired t-test, t(58) p < 0.05). **(f)** Effect of the time elapsed between the previous DEV or the previous Up state on subsequent Up state evocation. Violin plots from an example animal show trials which evoked (green) or failed to evoke (pink) Up states, and the time since a previous Up state (left) or since previous DEV (right) (* p < 0.05, Wilcoxon test,). **(g)** Violin plots for comparison of time since previous Up state for evoked (green) and spontaneous (black) Up states, from the example animal (* p < 0.05, Wilcoxon test). (h) Cumulative distribution function for time since previous Up states, for evoked and spontaneous Up states. **(d,e,i,j,k)** As in **b**, **c**, **f, g, h** but mean responses for each animal were calculated before these population analyses.

We then compared responses to DEV and STD stimuli of different frequencies in AC-L (Figure 3c,e) to assess whether standard tones could also evoke Up states. Because cortical neurons are not sensitive to thalamic inputs following an Up state (Compte et al., 2003; Watson et al., 2008), we hypothesised that if DEVs trigger Up states by thalamic input, they would be less likely to be evoked following a more recent Up state. To this end, we tested whether the time since the previous Up state influenced the probability of putative Up state initiation. A two-sided Wilcoxon rank sum revealed that in our example animal deviants that failed to evoke an Up state had a more recent prior Up state, consistent with a refractory period (Z = 5.2033, p < 0.001 Figure 3f). This effect was also present in the population animal data (Z = 8.0699, p < 0.0001 Figure 3i), supporting thalamocortical triggering of Up states. In contrast, there was no effect on whether a deviant would evoke an Up state based on the time since the previous deviant stimulus. To further confirm our results are not due to entrainment, we compared the time since a previous Up state for spontaneous and evoked Up states (Figure 3g,j). Deviant-evoked Up states occurred more recently following a previous Up state, indicative of triggering Up states by DEVs and not simple entrainment. Further, a Cox proportional hazards regression confirmed that auditory deviants increase Up initiation probability above that expected from inter-Up state timings for spontaneous Up states (Figure 3k; p =0.0313; β=.0427). These data indicate that Up state history may determines Up state induction more that stimulus timing history, providing evidence for Up states being evoked and not neural entrainment to deviants.

If spontaneous and evoked Up states are similar event, they should have similar dynamics. To test this we divided the data into 3 categories: 1) Up phases that were initiated by a DEV, 2) spontaneous Up phases, and 3) a control condition of DEV responses when an Up state was not evoked (Figure S1). Comparison of these three responses revealed similar response shapes for Up phases that occurred spontaneously and those evoked by a DEV. In comparison, no responses were observed when the DEV failed to evoke an Up phase, consistent with the lack of response in Figure 2d,g.

To test the hypothesis that Up states are evoked by positive prediction errors, and not simply by release from adaptation or probability, we compared deviant responses to those of the many-standards (MS) control. The many standards control presents random tones, including the 10 and 14kHz tones used in the oddball (Figure 4a). Deviant-evoked responses from the left auditory cortex showed much larger, sustained responses to auditory deviants and not the many-standards control (Figure 4b). Amplitudes were compared for the AC-L within a time window of 100 to 300 ms, using a linear mixed-effect model with DEV/MS as a fixed effect and Animal, Trial and tone frequency as random effects. This confirmed larger responses to deviant stimuli than many-standards stimuli (ANOVA; F(1, 798) = 96.16, p < 0.001). For the PFC-L, amplitudes were compared within a time window of 200 to 600 ms, and also confirmed larger DEV-evoked responses (ANOVA; F(1, 158) = 26.44, p < 0.001).

**FIGURE 4.**
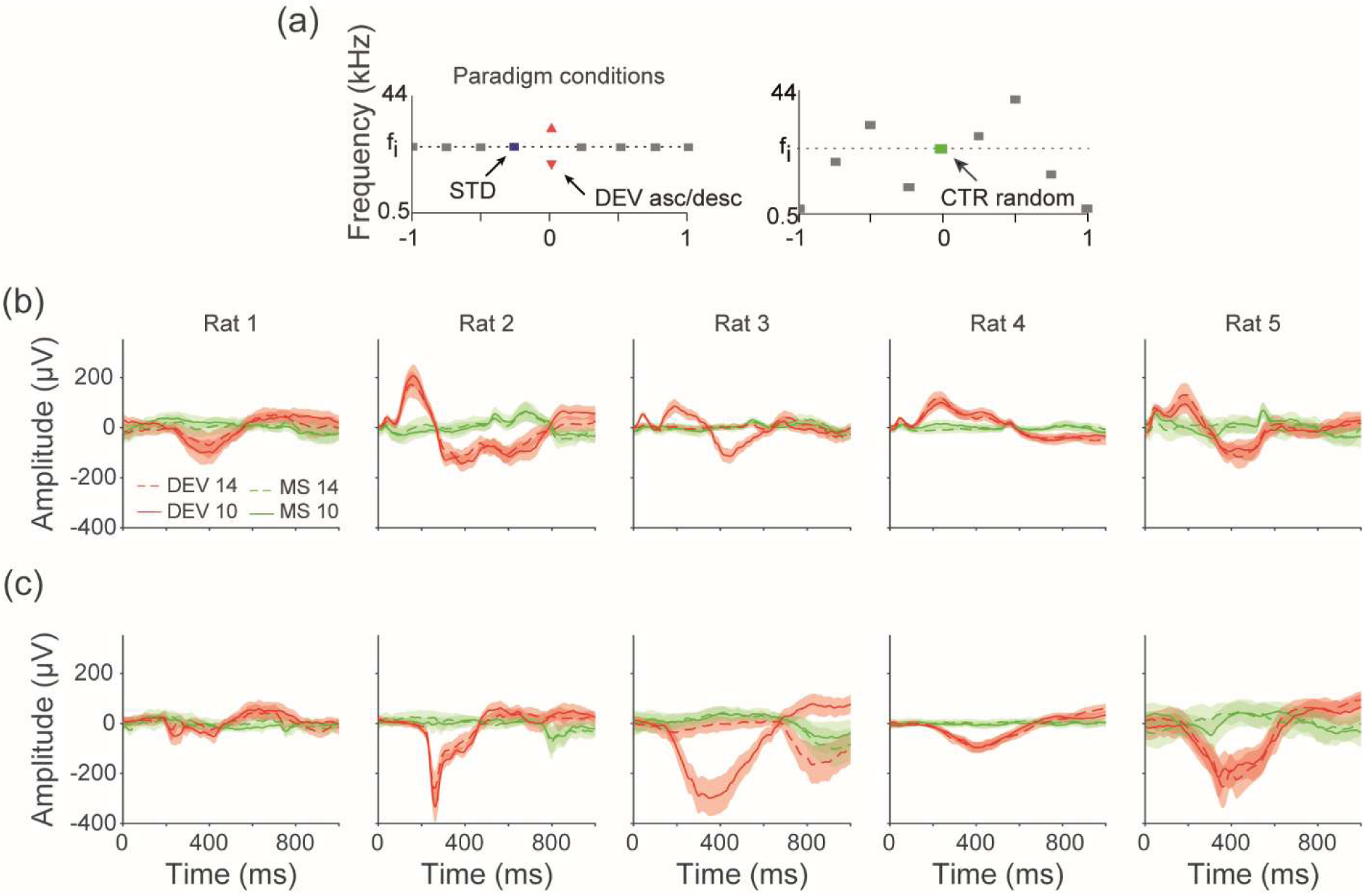
**(a)** We used auditory stimuli which consisted of a sequence of standard (STD) tones (blue) followed by a deviant (DEV) tone (red) of increasing or decreasing frequency placed randomly with a 10% probability. The paradigm allowed analysis of responses to the same tone but in a variety of contexts. We used a many-standards control (CTR). Mean ± 95% CI plots of Deviant (red) and Many-standard response for the AC-L **(a)** and PFC-L **(b)**. The dashed lines indicate the DEV and many-standard at 14 KHz, and the solid lines indicate those at 10 kHz. Mean ± 95% CI.

### Does anaesthetic depth modulate deviant-evoked cortical Up states

We hypothesised that anaesthesia is necessary for deviant-evoked Up states. To preliminarily test this, we chronically implanted an ECoG array in one rat and recorded activity 11 weeks post-implant. We compared responses in the awake state and under urethane anaesthesia in the same animal, using identical auditory stimuli one hour after anaesthetic induction. At this point, the animal reached a surgical level of anaesthesia and showed areflexia. We presented a flip-flop oddball paradigm (75 ms duration, 250 ms SOA; 10% DEV probability; mean inter-DEV interval 2.5 s; 10/14.142 kHz tones at 75 dB SPL;). During wakefulness, LFP traces showed smaller amplitudes (Figure 5), indicating a desynchronised state. One hour after urethane administration, the brain entered a synchronised state with clear cortical slow oscillations (Figure 5a). In both the PFC-L (Figure 5b) and AC-L (Figure 5c), deviant stimuli evoked stronger long-latency responses under anaesthesia than in the awake state, consistent with cortical Up states.

**FIGURE 5.**
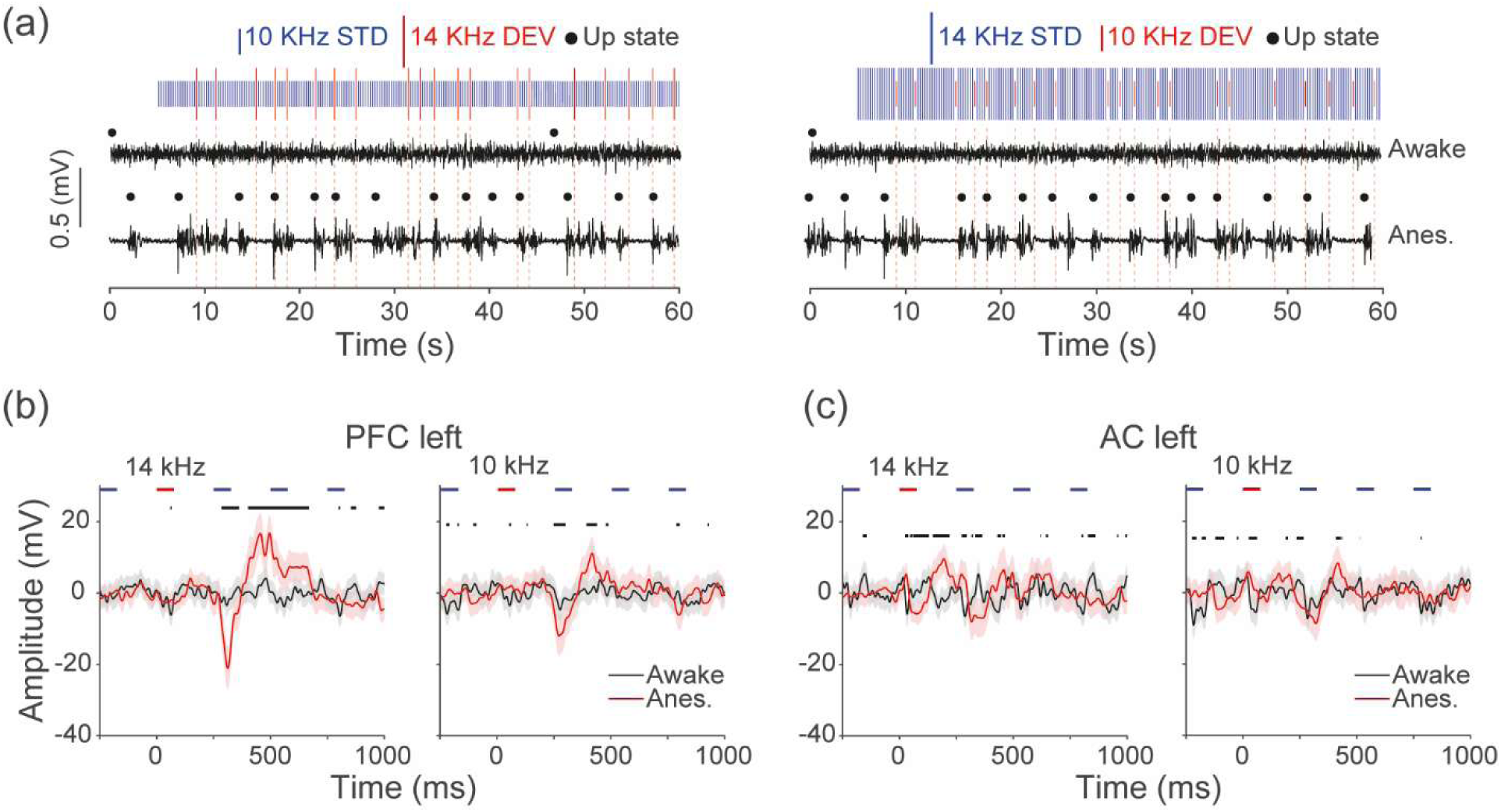
Up states may occur only under anaesthesia. **(a)** We compared DEV responses in awake and under anaesthesia to study the presence of Up states. LFP trace examples are from awake and 1 hour after urethane injection. A flip-flop oddball paradigm was presented with a 250 SOA and 10% DEV probability. Stimuli length was 1600, 480 DEV, 43 minutes. **(b**) LFP plot from mPFC-L for DEVs of 14 KHz (left) and 10 KHz (right), black line represents the awake recording and red the anaesthetised one. On x axis 0 represents the DEV onset. Mean ± 95% CI. Black bars show significance (two sample unpaired t-test, p < 0.05). **C(c)** The same as B but for data from the AC-L.

To also preliminarily assess how anaesthetic depth modulates long-latency deviant responses, we administered incremental doses of urethane to one rat. We presented a classical oddball paradigm (same parameters as above) and used a fixed 0.1 mV threshold to identify Up states across time to allow direct comparison. We began recording 6 hours after initial induction (baseline) and repeated stimulus presentations every 15 minutes over a 6-hour period while administering supplemental urethane. LFP traces from 14 kHz and 10 kHz deviants (Figure 6a) showed a progressive decrease in Up state frequency with increasing anaesthetic depth. Synchrony between deviant onset and Up state initiation increased with each urethane dose until very deep anaesthesia reduced the likelihood of exiting Down states (Figure 6b). We then calculated the percentage of deviants that evoked an Up state, which increased in parallel with synchrony as urethane levels rose (Figure 6c). These preliminary findings indicate that both the presence and depth of anaesthesia could influence the probability and timing of deviant-evoked Up states, though further studies would be required to confirm this.

**FIGURE 6:**
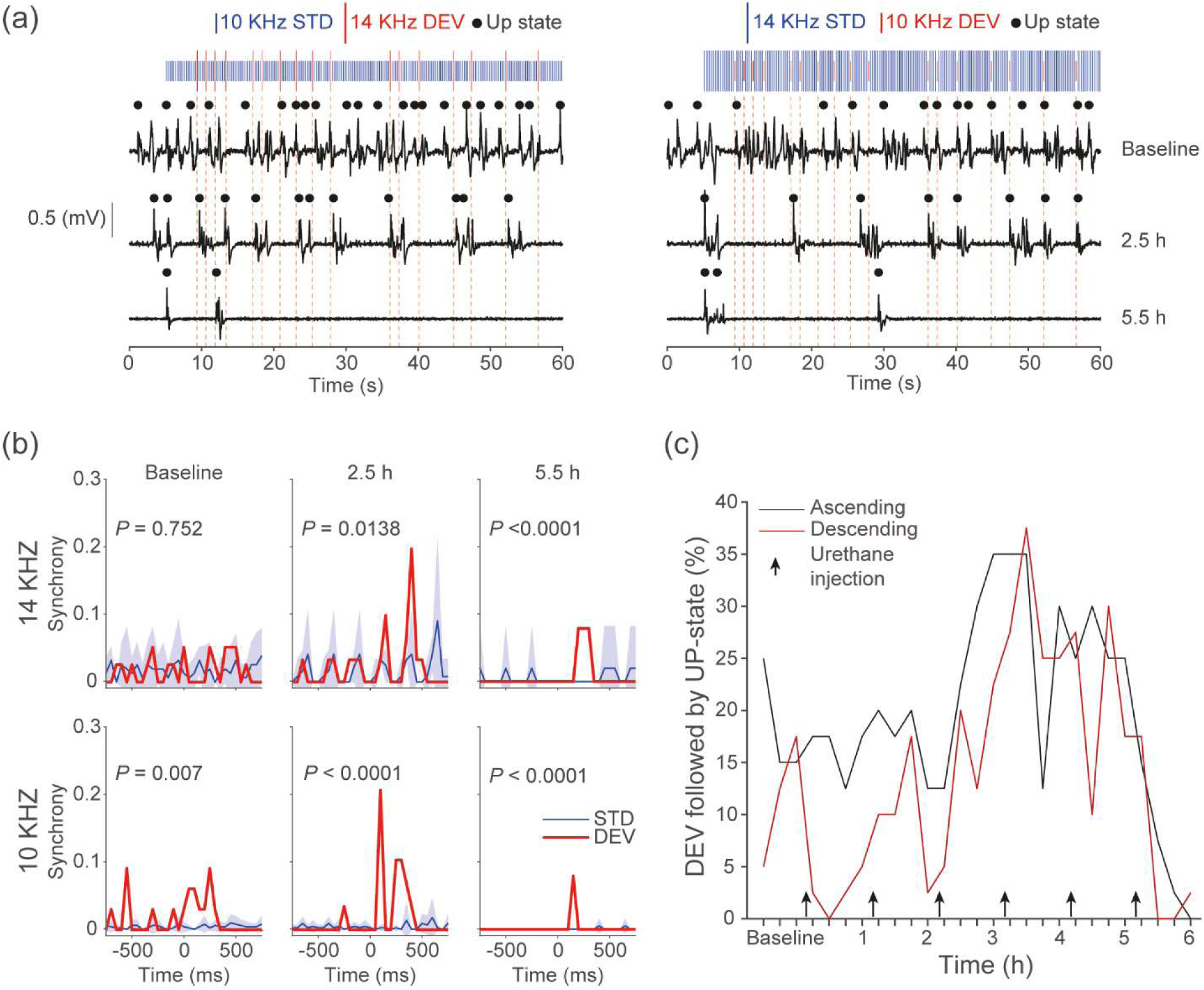
Deviants evoking cortical Up states may be dependent on anaesthetic depth. **(a)** In one animal, we gave supplemental urethane anaesthetic doses to assess the impact on Up state initiations. LFP traces from baseline, is 6 hours after initial urethane induction; 2.5 and 5.5 hours after 5 and 8 supplementary urethane dosages; for an example animal. A classic oddball paradigm was presented with a 250 SOA and 10% DEV probability, to allow presentation every 15 minutes. Data shown are the initial 60 seconds, for clarity. Stimuli length was 75 ms, 40 DEV, 11 minutes. **(b)** Cross-correlation synchrony between Up state initiation and DEV or STD stimuli for ascending or descending oddballs (top row & bottom row, respectively). Two sample unpaired t-test, t(15)). Mean ± 95% CI. **(c)** Percentage of DEV that are followed by Up states, throughout further urethane administration, for ascending or descending oddballs (black & red, respectively).

## DISCUSSION

Our ECoG recordings in rats revealed spontaneously occurring Up and Down states under urethane anaesthesia. When we presented an auditory oddball paradigm, only DEV stimuli evoked all-or-nothing long-latency responses that closely resembled features of cortical slow oscillation Up states. These possible deviant-evoked responses emerged first in AC at the shortest latency and then spread to other regions, consistent with the traveling wave dynamics of Up state propagation (Muller et al., 2018). Deviants trigger Up states more reliably when a longer period had elapsed since the previous Up state, consistent with a refractory period. Standard and many-standard stimuli failed to initiate Up states. Our preliminary data suggest that only anaesthesia enables deviant-evoked Up states, and the effect scales with anaesthetic depth, though these findings are based n individual animals. When deviants did not trigger an Up state, we observed minimal deviant-related activity, thus it is tempting to speculate that Up state initiation represents the principal cortical response to auditory deviance under urethane anaesthesia (Figure 3). These responses featured an early negative peak around 50 ms and 20 µV, consistent with the rat MMN (Jung et al., 2013b), followed by a later, larger peak at 200 ms and 200 µV that we interpret as Up state initiation, given its characteristics under anaesthesia. Our preliminary data suggest that only anaesthesia enables deviant-evoked Up states, and the effect scales with anaesthetic depth, though these findings are based in individual animals.

Although our study does not directly address the mechanism by which deviants evoke Up states, previous work has shown that both thalamocortical activity (Contreras & Steriade, 1995; David et al., 2013; Lemieux et al., 2014; Rigas & Castro-Alamancos, 2007) and sensory stimuli (Hasenstaub et al., 2007; Ji & Wilson, 2007; Petersen et al., 2003) can trigger Up states. Neurons in the auditory thalamus exhibit stimulus-specific adaptation and respond strongly to deviant stimuli (Antunes et al., 2010; Parras et al., 2017; Valerio et al., 2024). Based on this, we suggest that enhanced deviant responses in the auditory thalamus during the oddball paradigm is likely to increase thalamocortical drive and thereby evoke cortical Up states. Our findings support this interpretation, as Up states appear to initiate in the AC-L, the primary site of thalamocortical input during right-ear stimulus presentation. These Up states then propagate across the cortex as traveling waves at an estimated rate of ∼30 mm/s (Ruiz-Mejias et al., 2011; Stroh et al., 2013), which likely accounts for the delayed responses observed in AC-R and mPFC-L (Figure 3b,d). Future studies using this same paradigm while selectively inhibiting thalamocortical input could confirm this proposed mechanism.

If the mechanism we suggest is correct, adaptive and predictive processes that shape thalamocortical activity may play a central role in generating sensory cortex responses to deviants during unconscious states. Thalamocortical input would evoke cortical Up states from the primary sensory cortex and propagate them across the cortex as traveling waves (Maria V. Sanchez-Vives & David A. McCormick, 2000; Massimini et al., 2004; Muller et al., 2018). An enhanced thalamic response to deviant (but not standard) stimuli could therefore generate a long-latency mismatch response throughout the cortex. Although such responses may reflect stimulus-specific adaptation or prediction error within the thalamus, they may not represent true prediction error signals in all cortical areas. Instead, these responses may arise solely from increased thalamocortical drive sufficient to initiate an Up state. Future studies that record cortical Up states while manipulating thalamocortical pathways could evaluate this potential mechanism. If this account holds, interpreting long-latency prediction error signals in the cortex during unconscious states requires caution.

We also tested whether omitted stimuli could evoke cortical Up states, using two omission paradigms with different SOAs. In one condition (Rats 3 and 4), we used a 250 ms SOA (75 ms duration 6.45% omission probability; inter-omission interval 3–4.75 s; STD 10 kHz, 60 dB SPL), and in the other (Rats 5 and 6), we used a 125 ms SOA (6.45% omission probability; inter-omission interval 1.5–2.375 s; same tone parameters). In all 4 animal, we observed no evidence that omissions alone initiated cortical Up states (Fig. 7). However, in the 250 ms SOA condition, the post-omission stimuli (POST) frequently evoked Up states, and the timing of these responses corresponded more closely with the onset of the post-omitted tone than with the omission itself (two-sample t-test, *p* = 0.0019). These findings remain preliminary, and future experiments with larger samples and optimiszed paradigms will be necessary to clarify whether omissions can modulate Up state dynamics under anaesthesia. Omissions induce negative prediction errors, which are present in a lower number of neurons and at lower activity rates than positive prediction errors or responses to standard stimuli (Auksztulewicz et al., 2023; Awwad et al., 2023; Lao-Rodríguez et al., 2023a). Negative prediction errors would therefore be unlikely to increase cortical activity sufficiently to evoke Up states. POST responses are greater than omission resppnses in single neurons of the auditory cortex of anaesthetised rat (Lao-Rodríguez et al., 2023b), therefore POST stimuli would be more likely than omissions to evoke Up states. In future studies, an alternative paradigm such as the duration oddball could be used, which also produces prediction errors and we would hypothesise this to also evoke Up states.

**Figure 7:**
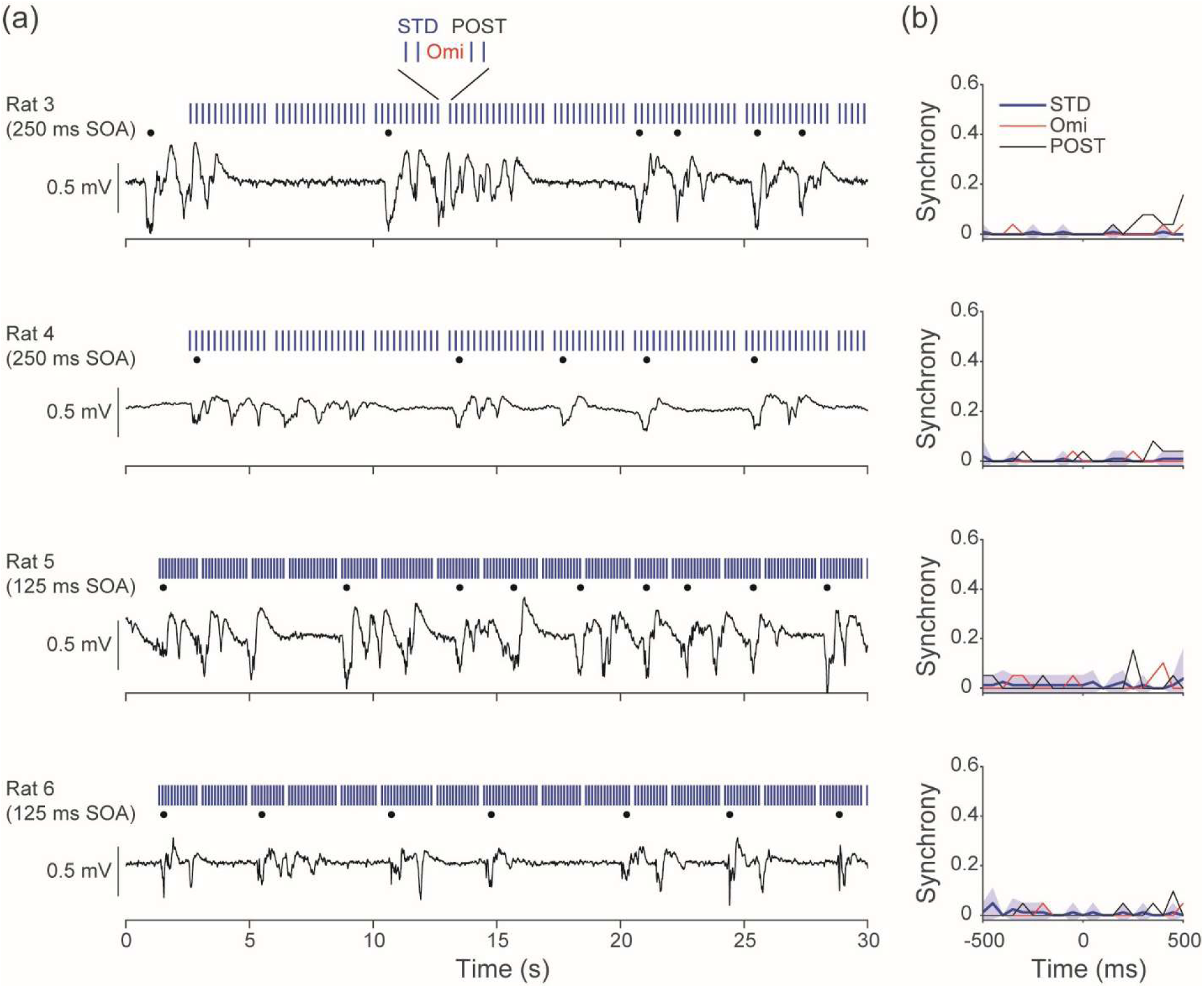
Omissions do not evoke cortical Up states. **A)** LFP traces from the mPFC-L. We tested in animals 3-6 (rows), omission paradigms consisting of a repeating STD (blue bars) and omission (spaces) for Up state initiation (black dots). We tested animals 3&4 with a 250 ms SOA, and animals 5&6 with a 125 ms SOA. For clarity only the initial 30s are shown, full stimuli length was 620 stimuli, 40 DEVs, 77/155s. **B**) Mean ± 95% CI plots of cross-correlation synchrony between Up state initiation and omission or STD stimuli presentation revealed that silent omission or repeating STD stimuli did not initiate cortical Up states in any animal. However, the post-omission stimuli were able to evoke Up states in the 250 ms SOA condition in Rats 3 & 4 (two-sample t-test, p = 0.0019). The latency of Up state evoking is consistent with Up state evoking by the post-omission.

Previous studies have described long-latency cortical responses to auditory deviants during anaesthesia and synchronised sleep, but have not linked these responses to cortical state changes (Casado-Román et al., 2020; Hockley & Malmierca, 2024; Loewy et al., 2000; O’Brien, 1982; Parras et al., 2017; Tóth et al., 2024; Xiong et al., 2024). Based on our present data, we suggest that long-latency deviant responses may reflect a state change within the desynchronised brain state of deep anaesthesia, which resembles non-rapid eye movement (NREM) sleep (Harris & Thiele, 2011). MMN is a sensory-evoked potential that serves as a biomarker for numerous neuropsychological disorders (Kocagoncu et al., 2021; Lee et al., 2017; Sapey-Triomphe et al., 2023; Sterzer et al., 2018), although debate continues regarding its underlying neural mechanisms (Fitzgerald & Todd, 2020; Garrido et al., 2009; Malmierca et al., 2019; Ross & Hamm, 2024). Studies have observed MMN and related neural activity during sleep (Nashida et al., 2000; Sallinen et al., 1994; Strauss et al., 2015) and under anaesthesia (Ayala & Malmierca, 2015; Koelsch et al., 2006; Malmierca et al., 2009; Quaedflieg et al., 2014), and some have reported that neuronal correlates of MMN, such as stimulus-specific adaptation, can even increase in certain anaesthetic states (Polterovich et al., 2018; Taaseh et al., 2011).

Our data show a typical rat MMN response at ∼50 ms in primary auditory cortex (Jung et al., 2013b), followed by a long-latency response at ∼200 ms that occurs exclusively in response to deviant stimuli. We attribute this later response to the initiation of cortical Up states, as it matches spontaneous Up states in waveform, occurs in an all-or-nothing fashion, propagates globally across the cortex as a traveling wave, and varies with anaesthetic depth. This long-latency activity may resemble the human P300 component of the event-related potential, which reflects novelty detection, context updating, and higher-order cognitive processing (Polich, 2003; Polich & Kok, 1995; Verleger, 2020).

By administering supplemental doses of urethane, we demonstrated that the ability of deviants to possible trigger Up states depends on anaesthetic depth (Figure 6). When we compared this to recordings from an awake animal, we found that Up state responses did not occur globally during wakefulness (Figure 5). Anaesthetic depth could influences the duration of Up and Down states, with deeper anaesthesia increasing the time spent in the Down state (Mondino et al., 2024b). At low urethane doses, some studies report increased cortical complexity (Dasilva et al., 2021; Torao-Angosto et al., 2021), while others show that animals spend more time in NREM-like states than in REM-like states (Mondino et al., 2024b). In our data, animals in a more synchronised state (characterised by robust and rhythmic Up and Down states) showed more consistent Up state initiation in response to deviants (Figure 6). This dependence on anaesthetic depth likely reflects the increased difficulty of transitioning out of the Down state under deeper anaesthesia (Mondino et al., 2024b).

The long-latency deviant response under anaesthesia may be the result from a state change. However, the relationship between processing in the synchronised brain and awake brain remains unknown. We did not observe any long-latency state changes during wakefulness, which may reflect the more subtle and localised cortical state dynamics typical of wakefulness, or the passive nature of our awake recordings, which lacked directed attention to the deviant (Harris & Thiele, 2011; Speed et al., 2019). Because the Up state response follows the traditional MMN, the relevance to awake processing may be or activation of cortical ensembles (Hamm et al., 2021) or behavioural decisions and state transitions (Cole et al., 2024b; Gong et al., 2024) though further studies would be required to confirm these hypotheses. Traveling waves as in Up states have also been proposed to underlie predictive processing, with forward or backward travelling strength dependent on the weighting of priors to inference (Alamia & VanRullen, 2019), so the forward travelling waves observed here may provide evidence for this model.

Our study suffers several limitations. First, the definition and detection of Up states based on LFPs may conflate stimulus-evoked potentials with genuine slow-oscillations transitions. We have tried to reduce this possibility by detecting Up states based on large long-latency peaks not expected to be regular auditory-evoked potentials. Second, the small number of animals for the preliminary study on how anaesthetic depth alters Up state initiations restricts reproducibility and generalizability. Finally, while the findings are intriguing, extrapolations to human attention, P300 or disease models are premature given that most data were obtained under deep anaesthesia. Future studies with larger samples should confirm or deny if the present hypothesis is due to anaesthetic depth, and its relevance to desynchronised brain states.

## Author Contributions

**Laura H Bohorquez:** Methodology, Software, Formal analysis, Investigation, Writing - Original Draft, Writing – Review & Editing, Visualization

**Manuel S Malmierca**: Conceptualization, Writing – Review & Editing, Supervision, Funding acquisition

**Adam Hockley**: Conceptualization, Methodology, Software, Formal analysis, Investigation, Writing - Original Draft, Writing – Review & Editing, Visualization, Supervision, Funding acquisition

## Data availability statement

- All data has been deposited at Zenodo at 10.5281/zenodo.15791248
- Analysis code has been deposited at Zenodo at 10.5281/zenodo.15791285

### ABBREVIATIONS

ABR: Auditory Brainstem Response
AC-L: Auditory Cortex Left
AC-R: Auditory Cortex Right
DEV: Deviant
ECoG: Electrocorticography
LFP: Local Field Potential
MMN: Mismatch Negativity
mPFC-L: medial Prefrontal Cortex Left
mPFC-R: medial Prefrontal Cortex Right
NREM: Non-Rapid Eye Movement
POST: Post-deviant stimulus
RMS: Root-Mean-Square
SOA: Stimulus Onset Asynchrony
STD: Standard
TDT: Tucker-Davis Technologies

## SUPPLEMENTAL INFORMATION

**Figure S1.**
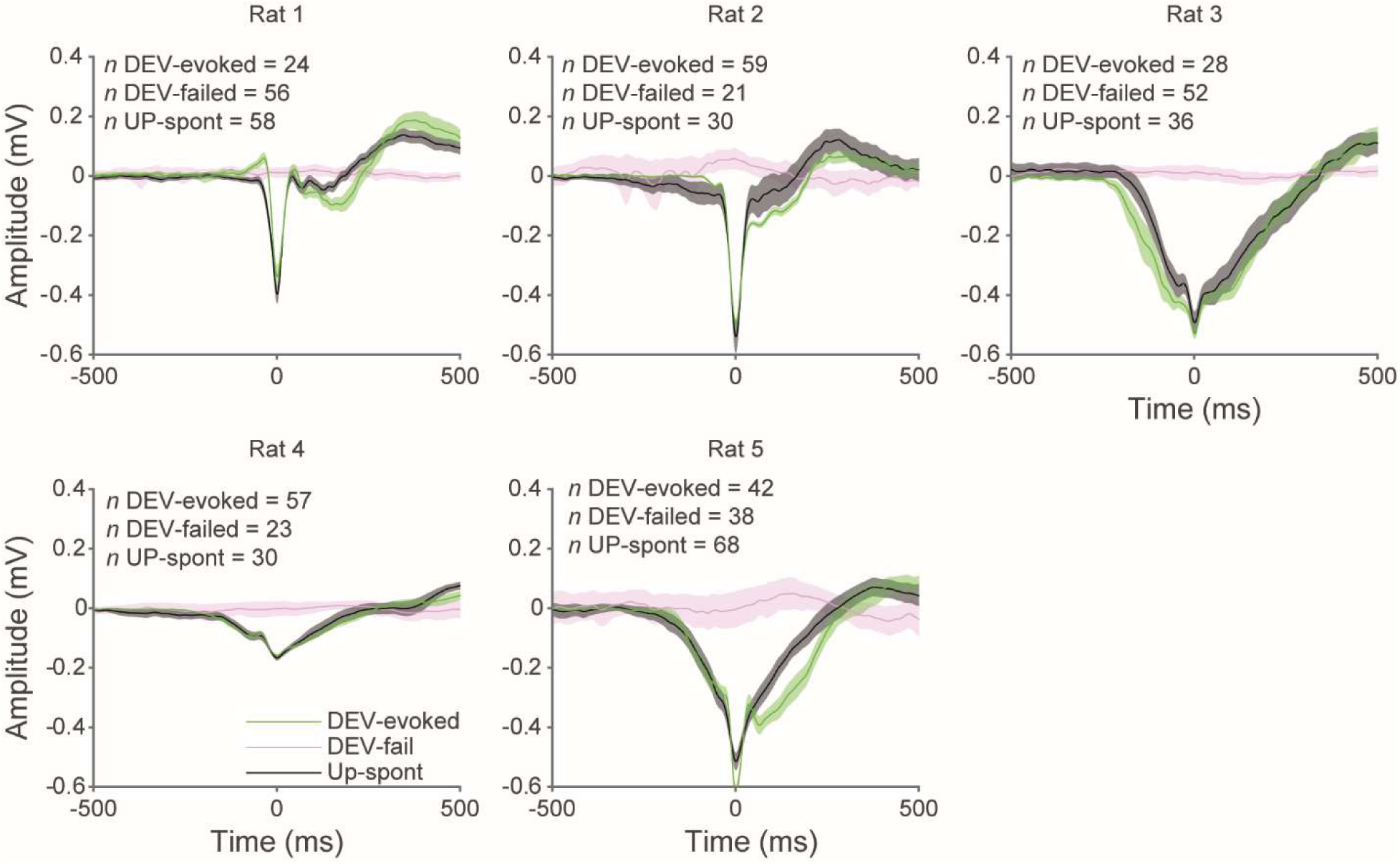
DEV-evoked and spontaneous Up states have similar dynamics Mean ± 95% CI plots of Deviant-evoked Up states (green) and spontaneous Up states (black) show similar response profiles. In comparison, when deviants failed (pink) to evoke an Up state, no other response is observed. Data from mPFC left electrodes, temporally aligned relative to negative peaks to allow shape comparison.

**Figure S2.**
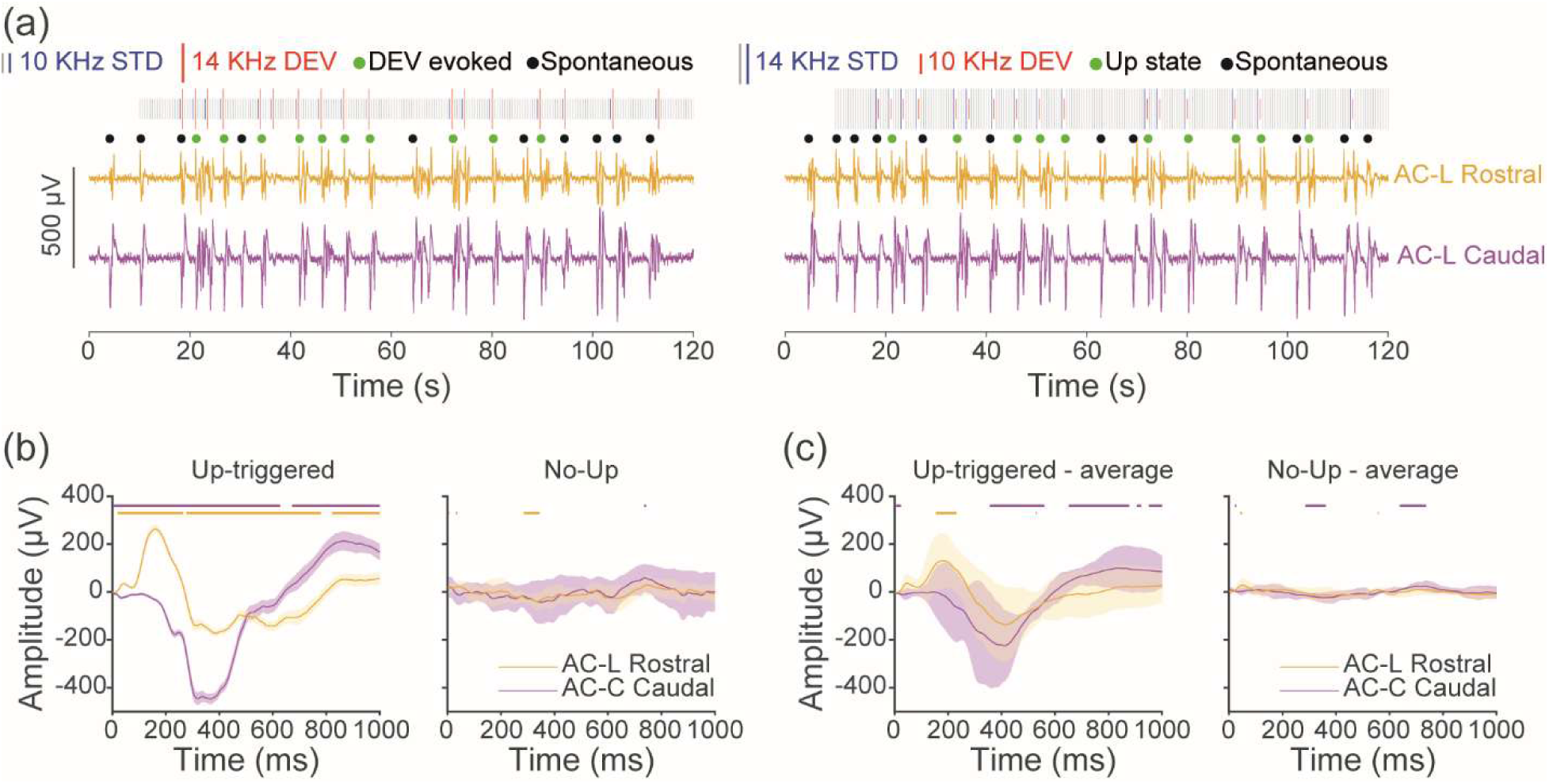
- Comparison of rostral and caudal AC recordings. **A**) LFP traces from AC-L Rostral (yellow) and AC-L Caudal (magenta) electrodes for an example animal. For each animal, a classical oddball paradigm was presented with a 500 ms SOA and 10% DEV probability. Stimuli length was 75 ms, 10% DEV probability, 82 minutes. Up state initiations were identified and classified as either evoked (AC-L Rostral response occurring 100-500ms after a DEV; green dots) or spontaneous (black dots). **B)** Mean ± 95% CI plots of DEV response from an example animal at different electrodes sites when the DEV is followed by an Up state (left) or not followed by an Up state (right). Yellow and magenta bars indicate the presence of significance from 0 for AC-L Rostral and AC-L caudal, respectively (one sample t-test, t(42) p<0.05).

